# Tunable Transcription Factor Library for Robust Quantification of Gene Expression Dynamics in *E. coli*

**DOI:** 10.1101/2021.11.16.468742

**Authors:** Vinuselvi Parisutham, Shivani Chhabra, Md Zulfikar Ali, Robert C. Brewster

## Abstract

Predicting the quantitative regulatory function of a TF based on factors such as binding sequence, binding location and promoter type is not possible. The interconnected nature of gene networks and the difficulty in tuning individual TF concentrations makes the isolated study of TF function challenging. Here we present a library of *E. coli* strains designed to allow for precise control of the concentration of individual TFs enabling the study of the role of TF concentration on physiology and regulation. We demonstrate the usefulness of this resource by measuring the regulatory function of the zinc responsive TF, ZntR and the paralogous TF pair, GalR/GalS. For ZntR, we find that zinc alters ZntR regulatory function in a way that enables activation of the regulated gene to be robust with respect to ZntR concentration. For GalR and GalS, we are able to demonstrate that these parlogous TFs have fundamentally distinct regulatory roles beyond differences in binding affinity.

## Introduction

Transcription Factors (TFs) are an important set of proteins that play a major role in controlling condition-specific cellular decision making. Techniques such as DNaseI footprinting^1^, SELEX^2^, ChIP-seq^3,4^ and their variants have enabled high resolution base-pair mapping of where TFs bind and which genes they control. However, predicting the direct regulatory effect of any given TF on a gene under its control, remains challenging; the ability to build genetic circuits from natural TFs or foretell the regulation of promoters directly from its architecture is still completely lacking. One challenge to these predictions is the interconnected nature of regulatory networks. Individual TF genes typically regulate (and are regulated by) several to dozens of different genes and so controlling the concentration of a TF systematically and thus the quantitative regulatory function of that TF at a target is convolved with network and “context dependent” effects that hide the direct role of the TF on the gene. As a result, predicting the quantitative input-output relationship between TF concentration and output of a gene based on regulatory architecture, *i*.*e*. the location, identity and sequence of the TF binding sites that contribute to a promoters’ regulation, is mostly not possible. However, tremendous progress has been achieved towards the predictive design of gene circuits and network architectures using model TFs^5–9^, although the toolbox of well characterized TFs is relatively sparse. Clearly, the characterization of a greater set of TFs would enable enhanced utility for biological engineering purposes while also deepening our understanding of why natural regulatory elements are built the way they are.

Here, we report the construction of a titratable copy of each of the 194 TFs in *E. coli* for the purpose of characterizing TF function. In this library, the copy number of any TF is controllable by induction rather than through indirect changes to growth or nutrient sources. The single-cell TF level is also measurable due to a fusion of the TF with the mCherry fluorescent protein. Importantly, expression of the TF is isolated from the natural regulatory interactions that would limit or complicate copy number control. In addition, the titratable TF construct is stably integrated at a constant genetic locus in the chromosome to avoid any copy number difference. The ability to titrate TFs precisely enables a direct and quantitative measure of the role of a specific TF in regulation or physiology. Similar approaches with individual model TFs have enabled deep understanding of the input-output function of those specific TFs^10–16^. The resource introduced here enables studies of TFs as a whole with the same quantitative control typically dedicated to model TFs. When combined with systematically designed promoters, this library enables careful examination of the input-output relationship of regulation for any TF in simple regulatory architectures that can reveal the fundamental regulatory function of these TFs. Overall, the goal in designing this resource is to enable detailed studies to characterize regulatory function of TFs in a less biased way (Fig 1A).

**Figure 1.**
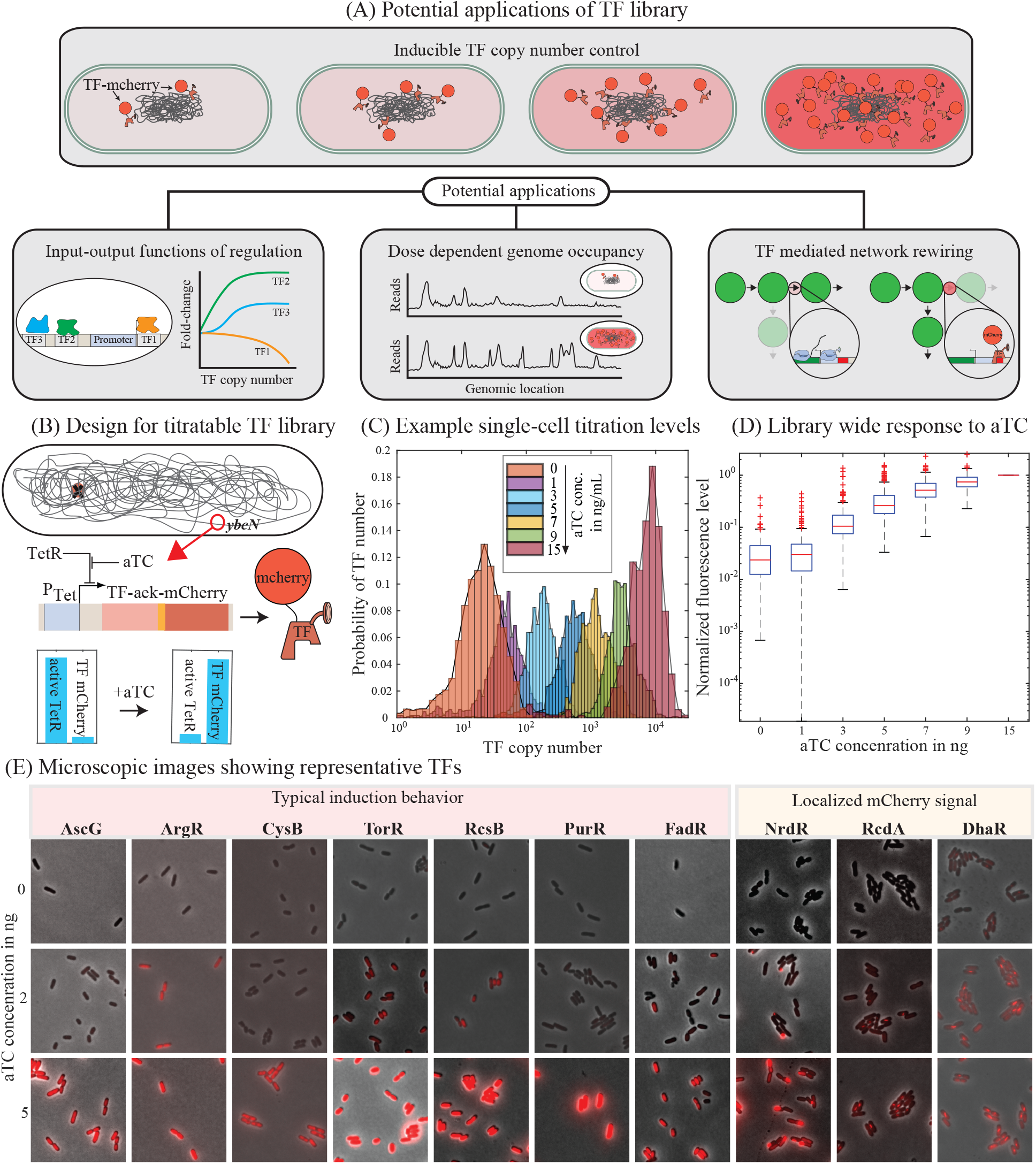
Design of titratable transcription factor library: **(A)** Potential applications of titratable TF library. **(B)** Schematic representation of the titratable transcription factor library strain. TF is deleted from its native locus and expressed from the *ybcN* locus as a mCherry-fusion construct and under the regulation of a tetracycline inducible promoter. **(C)** Histogram represents single cell levels of TFs for increasing concentration of the inducer, aTC. Representative TF used here is RcsB. **(D)** Box plot shows the distribution and mean mCherry levels expected for the given concentration of aTC for all the TFs in our library. **(E)** Representative microscopic images showing different features of the strains as result of TF titration. For instance, there is a significant change in length and mCherry signal for different levels of aTC in CysB library strains. DhaR, NrdR, and RcdA shows localized mCherry signal.

In the remainder of the manuscript we discuss the construction of the library followed by an account of the inducible TF copy number range of the library strains and the physiological impact of controlling the TF copy number in each strain. Finally, we give two examples of uses of the library to characterize TF function. In the first vignette, we examine the unique regulatory architecture of heavy-metal responsive TF, ZntR and in the second, we examine the regulatory differences in a pair of paralogous TFs, GalR and GalS. In both cases, we find that controlling TF number enables unique insight into the fundamental regulatory behavior of these two groups of TFs.

## Results

### Transcription Factor Library

Construction of the titratable TF library is described, in detail, in the Methods section and in Fig 1B. Briefly, each of the 194 TF gene is deleted from its native locus, and the corresponding tf-mcherry fusion gene (with an aek-linker sequence (AEAAAKEAAAKA) separating the TF and mCherry genes), is expressed from a tetracycline-inducible promoter and integrated at the *ybcN* locus. The inter-protein linker would likely not interfere with the bio-activity of the TF and enhance its stability^17^. In addition, *tetR* gene is integrated at the *gspI* locus and the constitutive gene product (TetR) will repress the expression of the TF until the inducer, anhydrous tetracycline (aTC) is present. Upon addition of aTC, the TF gene is expressed and is tracked by measuring the mCherry levels at a given aTC concentration (histogram in Fig 1C shows the single-cell level distribution of mCherry fluorescence for one representative TF, RcsB).

There are at least 198 genes of *E. coli* listed as transcription factors in RegulonDB^18^. Our titratable TF library consists of 194 pair (TF knockout and the corresponding titratable strain) of strains in total. The TFs included in our library can be classified into seven functional categories (Fig S1A): (1) transcriptional repressors (53 genes), (2) transcriptional activators (39 genes), (3) dual-regulators (74 genes), (4) histidine sensor kinase of the two-component system (18 genes), (5) DNA-binding regulator of the toxin/antitoxin system (8 genes) (6) multi-functional regulator (4 genes), and (7) pseudogenes (1 gene). There are at least two pseudo TF genes (*gatR* and *glpR*) in *E. coli* genome. Of these two pseudo genes, *gatR* is inactivated by an “IS element” inserted in the middle of the gene^19^, and as such it is not included in our library. On the other hand, for *glpR*, several genetic variants are reported in multiple rounds of sequencing. The *E. coli* MG1655 whole-genome sequence listed in NCBI has a single nucleotide insertion causing a frame-shift mutation. However, the *glpR* gene amplified from our lab stock of *E. coli* MG1655 does not contain this insertion (alleviating the frame shift), and hence that variant is included in our library. Despite repeated trials, construction of *relE*, toxin gene from a native toxin/antitoxin pair was unsuccessful. It is possible that even the “leaky” levels of RelE is enough to overwhelm the native expression of the toxin, RelB. As a result, *relE* and *relB* are also excluded from this library. Finally, the TF *alaS* is essential and the corresponding knockout is not available in the keio collection^20,21^ and, as such, we did not create a titratable strain for *alaS*.

Often times, altering the growth condition is used as a way to control TF copy number in *E. coli* via changes in gene dosage, protein dilution and network regulation^22^. However, these approaches do not always enable full control over a wide range of TF concentrations. For example, Fig S1G shows data from Schmidt *et. al*.^23^ measuring protein copy numbers over a wide range of growth conditions, it is clear that many TFs are not well controlled in this fashion and often times growth rate is a poor predictor of TF copy number. By expressing TFs from a controlled tetracycline-inducible promoter (at a common genetic locus), we eliminate many of the native transcriptional regulatory network features that otherwise influence the number of TFs and create a precise control of the TF copy number through induction (Fig S1H).

For 14 of the library strains, the fluorescence signal was very low in the plate reader, for these strains confirmation of the TF titer with aTC required single-cell wide-field microscopy. In 11 of these cases, the signal was either very low or the fluorescence signal was localized to specific regions of the cell (typically at the mid-cell or at the poles, see TFs: NrdR, RcdA and DhaR in Fig 1E). However, two TFs, CytR and RbsR (Fig S1E and F, dashed lines) did not show any titratable mCherry levels and the global regulator, Crp, was expressed at very high and constant levels despite of the inducer concentration. Re-sequencing of the modified regions in the library strains for Crp, CytR and RbsR did not show any mutation in the regions of interest. The constant mCherry signal for Crp could be because of some internal regulations within the coding region of the gene itself. Although many TFs have internal regulatory sites, we did not alter the natural coding sequence of any TFs in the library. For CytR and RbsR, the measured mCherry signal was very low and multiple factors such as alteration of the emission/excitation spectrum of mCherry due to the protein fusion could account for it. In the end, 191 TF strains in our library are titratable and show a similar pattern of titration saturating at around 3 ng/mL aTC (Fig 1D and S1H).

### Quantification of TF copy number

The mCherry fusion allows for the direct quantification of absolute TF copy number based on the proportionality between fluorescent signal (*I*) and the number of fluorophores (*N*), *I* = *νN*. The arbitrary fluorescence signal measured from a microscope or flow cytometer can be converted to the number of fluorescent proteins by estimating for the calibration factor, *ν*. The techniques to measure *ν* typically involve measuring either the fluorescence of a single molecule in photobleaching experiments^24–26^ or the measurement of a larger molecule that contains a fixed number of fluorescent protein^27,28^. An alternative method for quantifying TF copy number from fluorescence signal involves measuring fluctuations around the mean of a stochastic event that involves the fluorescent protein; two examples include measuring fluorescence level differences of two daughter cells immediately after division (Fig 2A)^7,29^ or measuring fluctuations in the fluorescent bleaching trajectory of fluorophores in a single-cell^30–32^. In our library strains, we used fluctuations in the partitioning of fluorescence signal between the daughter cells (*I*_1_ and *I*_2_) to measure the calibration factor for 10 strains (see Fig 2B and Materials and Methods “Estimation of calibration factor”) and hence the number of TFs. The estimate of *ν* is based on the assumption that the TF-mCherry proteins are randomly distributed between the two daughter cells upon division. In general, we find that the calibration factor for each TF is similar (Fig 2C) with one notable outlier (ArgR). The difference in calibration factor for ArgR could be due to higher order oligomeric states, or dependence of the co-factor arginine for DNA binding or a breakdown in the assumption of equal partitioning at cell division. Determining the calibration factor for all 194 TFs would be laborious and the actual value of *ν* depends heavily on experimental settings (microscope optics, exposure times *etc*), we chose to use the mean of the 10 measured calibration factors to estimate the number of TFs in each strain throughout the library. However, it is important to note that there may be cases where this estimate is significantly off, as is the case for ArgR. As such, the measurement of the TF number for other strains should be thought of as an estimate; if precise knowledge of TF numbers in a specific strain is required, an individual measurement in that specific library strain should be made. In Fig 2D, we show the response of the titratable library strains to aTC. We observed 100 − 1000 fold increase in TF numbers as aTC concentration is increased (Fig 2D). In Fig 2E, we compare the maximum induction level of TFs in our library to the measured TF numbers per cell, under 20 different growth conditions for *E. coli* from the work of Schmidt *et. al*.^23^. The vast majority of these data points are above one, indicating that our induction strains are capable of reaching to and beyond the physiological concentrations of most TFs.

**Figure 2.**
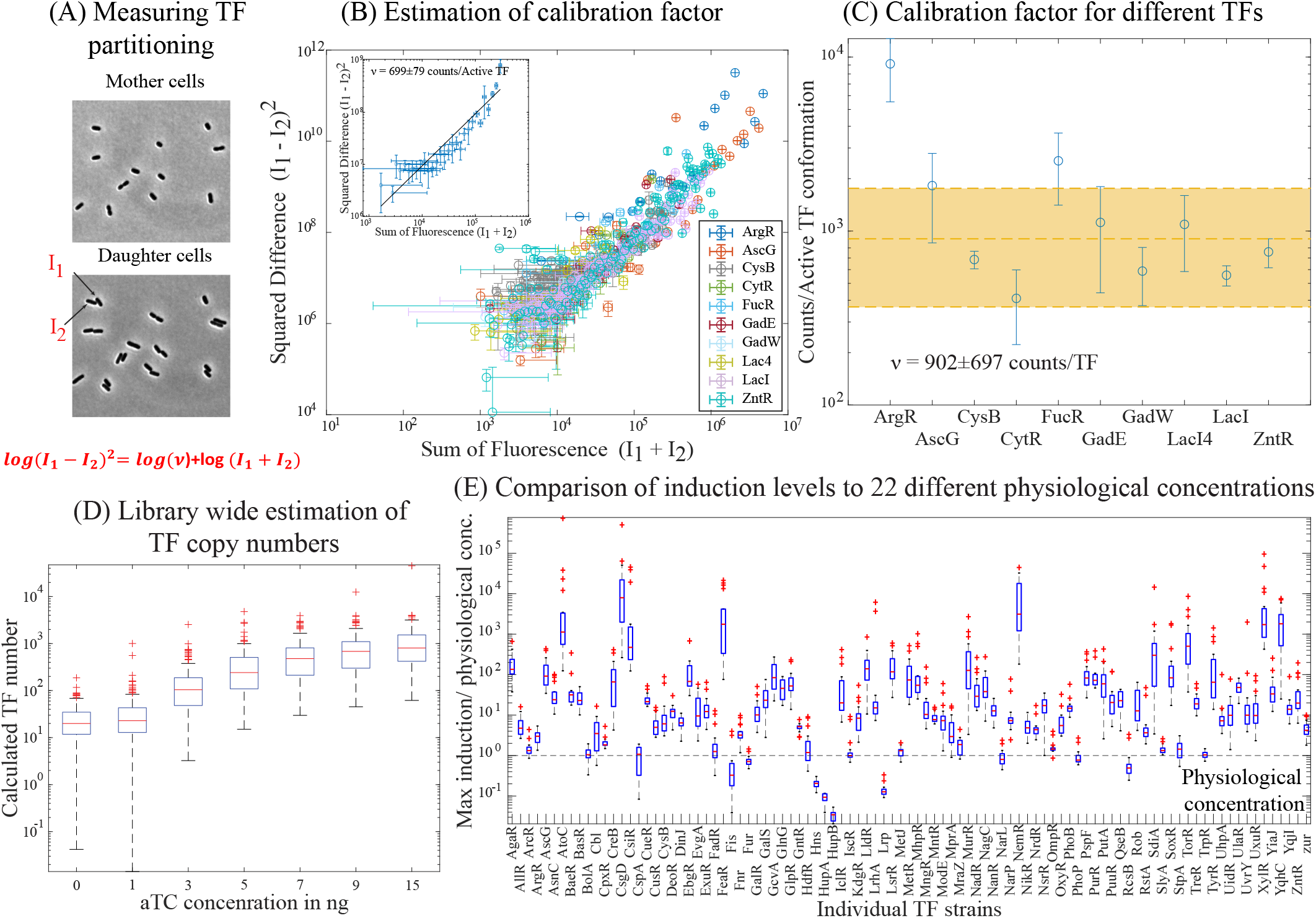
Estimations of TF number: **(A)** Representative images taken before (top) and after (bottom) one cell division in order to calculate the stochastic fluctuation in fluorescence distribution between the daughter cells, *I*_1_ and *I*_2_. **(B)** Plot showing the sum (*I*_1_ + *I*_2_) and squared difference ((*I*_1_ − *I*_2_)^2^) in fluorescence of the daughter cell pairs for different TFs. **(C)** Measured calibration factor for 10 different TF strains in our library. **(D)** The data in Fig 1D is converted to absolute TF numbers using the estimated mean calibration factor. **(E)** Comparison of the TF copy number at maximum induction to the measured protein copies/cell in the work by Schmidt *et*.*al*.^23^.

### Physiological effects of TF titration

TFs are directly (or indirectly) involved in rewiring the function of clusters of genes within the cell. Clearly, there will be physiological consequences for altering the concentration of some of the TFs. For instance, some TFs such as Crp, ArgR, CysB and MetJ are critical for essential metabolic pathways and under-producing or deleting these TFs may seriously affect the fitness of the corresponding strains under certain induction conditions. On the other hand, some TFs such as Nac and PdhR may be toxic when expressed at high concentrations^33^. Here, we will use growth rate in glucose minimal media as a proxy to evaluate the fitness or physiological effects due to the titration of each TF. The steady-state growth rate of each TF library strain is measured in different aTC concentrations and normalized to the growth rate of wild-type in the corresponding aTC concentration. Hierarchical cluster analysis results in 6 major clusters of growth phenotypes (Fig 3A). Furthermore, we calculate the correlation coefficient between growth rate and aTC concentration for each library strain (Fig 3B). The correlation coefficients are roughly tri-modal with one peak around negative correlation values, one near zero and the final less-defined peak for positive correlations. The majority of strains show a negative growth correlation with aTC. Importantly, this is not due to aTC toxicity; the grey shaded box in Fig 3B shows the correlation between wild type growth rate and aTC concentration. We also validate that there is no correlation between total mCherry levels and the growth rate (red shaded box, Fig 3). It is evident that the higher correlation between TF copy number and growth rates is primarily due to the physiological consequence of TF expression and not due to protein over-expression or aTC toxicity.

**Figure 3.**
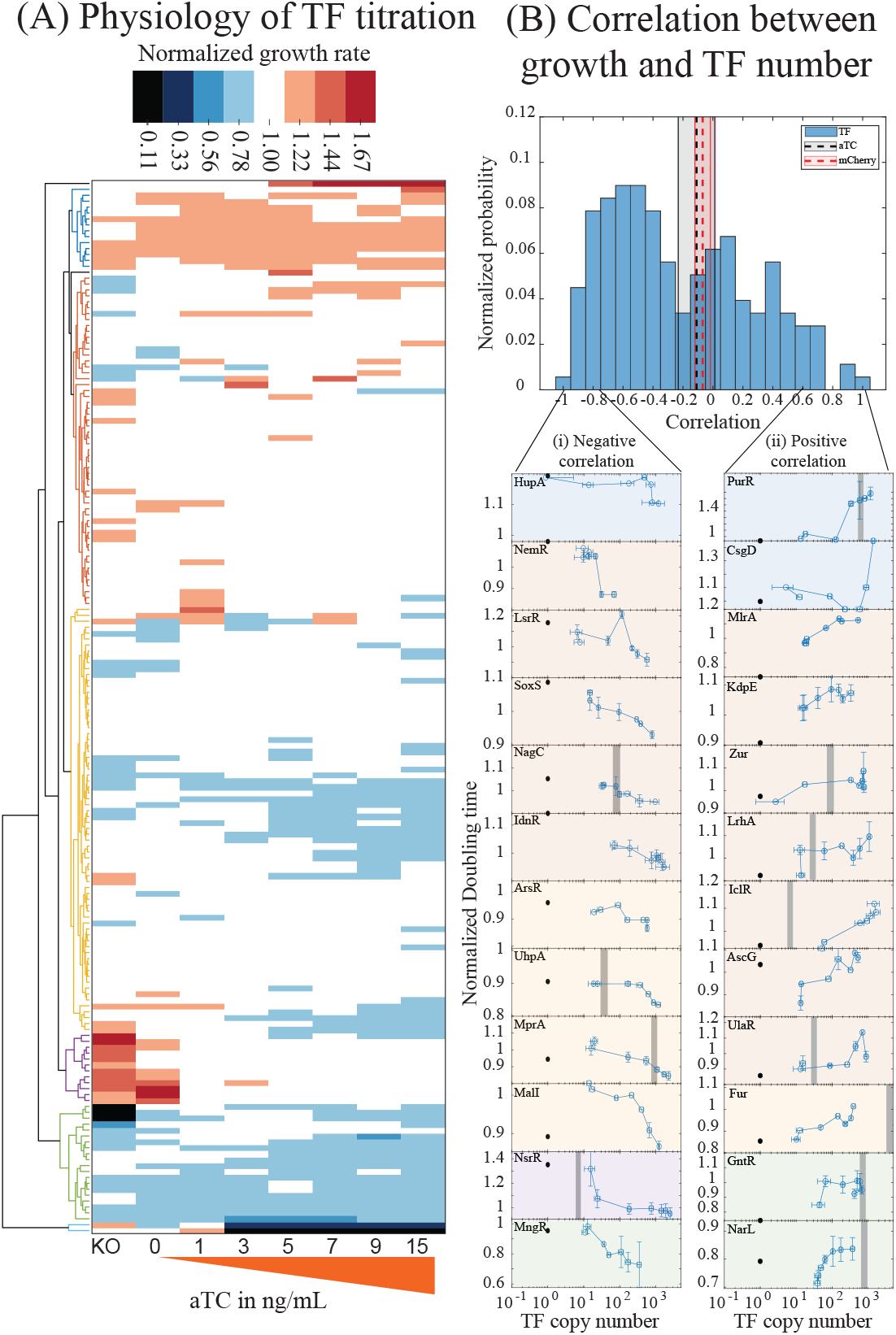
Physiology of the titratable library strain: **(A)** Cluster analysis of growth rates of the library strain in different concentration of aTC. The growth rate used here is normalized to the growth rate of wildtype MG1655 measured in similar aTC concentration. **(B)** Histogram showing the correlation between growth rates and the estimated TF copy number. Shown in grey shades is the correlation for wildtype MG1655 in different aTC concentration and shown in red shades is the correlation between growth rates and total mCherry. Lateral panel shows representative examples of strains showing high correlation between growth rates and TF copy number. The grey bar represents physiological TF copies/cell measured by Schmidt *et. al*.^23^. The panels are colored to match the grouping in the clustergram in Fig 2A

In Fig 3A, the top most cluster in blue corresponds to TF strains that grow faster than wild type at most aTC concentration. Despite growing faster than wild type, this cluster includes TFs that show both positive, and negative correlation between growth rates and TF copy number (Fig 3B, lateral panel (i): HupA and (ii): PurR and CsgD). PurR, a regulator of purine metabolism, exhibits the fastest growth rate with a doubling time of 41 ± 2 minutes (Fig 3B, lateral panel (ii): PurR). A few genes of this cluster including McbR, BluR and CsgD are involved in regulating biofilm formation. The second cluster, in red, has two bigger nodes. Top node include TFs whose knockouts grow slower than wild type and TF titration helps enhance the growth rate; some of which exhibit positive correlation between TF copy number and growth rate (Fig 3B, lateral panel (ii): MlrA, KdpE, Zur, LrhA, IclR, AsnC and UlaR). The Bottom node includes TFs with knockouts growing faster than wild type and TF titration retards the growth rate exhibiting a negative correlation between TF copy number and growth rate (Fig 3B, lateral panel (i): NemR, LysR, SoxS, NagC and IdnR). The third cluster includes TF strains that show overall reduced growth compared to wild type. The slowest doubling time in this cluster is 89 minutes for the TF, MntR involved in sensing heavy metal, manganese. In this cluster the growth rates across different aTC concentration is fairly constant however, there are exceptions (such as Fig 3B, lateral panel (i): ArsR, UlaR, MprA, MalI and lateral panel (ii): Fur) where we see a good correlation between growth rate and TF copy number. Fourth cluster in purple includes TFs with knockouts and the lowest aTC concentrations growing faster than the wild type. Further increase in aTC causes a reduction in growth rate (Fig 3B, lateral panel (i): NsrR). Strains with extreme growth defects are part of the green and cyan clusters. In the first node of the green cluster are TF strains where the knockout shows “no growth” or reduced growth and expression of just enough TF is sufficient to rescue the growth rate. All TFs of the amino acid metabolic pathway (ArgR, CysB, MetR, MetJ and LysR) belong to this cluster. In the second node are TF strains which show drastic decrease in growth rate upon TF titration. The slowest doubling time measured in this cluster is 115 minutes for the TF, BglJ. Cyan cluster has only two strains, Nac and PdhR, that stops growing beyond the 3 ng/mL of aTC in the medium. In summary, Nac and PdhR are the only two TF strains exhibiting severe growth defects hampering its use in the titratable TF expression.

### Case studies using TF titration library

In the following two sections we demonstrate how the TF titration library can be used to dissect the regulatory function of individual TFs. The library is particularly effective when combined with other genetic resources that allow for systematic control of promoter architectures. The vignettes of these section make use of two reporter libraries, the Zaslaver’s transcriptional reporter library^34^ and a “TF binding position library” from our lab which enables controlled movement of a TF binding site on a synthetic promoter^35^.

### Case study 1: Regulation by the zinc responsive TF, ZntR

The architecture of a promoter (*i*.*e*. the position, identity and specificity of TF and RNAP binding sites) is a fundamental indicator of overall regulatory activity of a gene. Elucidating the mechanisms of common promoter architectures can help lay ground rules to build well-defined genetic parts. For instance, the architecture of promoters involved in sensing heavy metals such as copper, zinc, gold and mercury are very similar across different bacterial species (Fig 4A). Interpreting the biophysical constraints of such promoter architectures will help in different biological applications such as in whole cell biosensors. The common promoter architecture of heavy metal responsive genes involve a single TF binding site acting as the spacer between the − 10 and − 35 promoter sequence. This architecture is common for MerR family of TFs in *E. coli* (such as the metal responsive TFs, *cueR* and *zntR*, responding to copper and zinc respectively) and is also found in TFs from other organisms such as *bltR* of *Bacillus* and *merR* of transposable elements^36^. Previous studies have examined this promoter architecture and determined that the mechanism of activation involves a distortion of the promoter DNA upon binding of the co-factor bound TF which can realign the -35 and -10 boxes that is separated by an unusually longer spacer of 19 − 20 bp (the optimal spacer length for *E. coli σ* ^70^ promoter is 17 bp^37,38^). These studies involve mutating or truncating the spacer sequence which in turn will abolish the TF binding site^36^. However, the precise input-output function of these types of promoters can often be difficult to tease apart due to their natural coupling to physiology and inherent feedback in regulation. We examined the zinc responsive TF, ZntR, as a model TF. The only known function of ZntR is in regulating the expression of ZntA, a transmembrane protein that mediates the export of zinc and other heavy metals. Less is known about the regulation of ZntR in *E. coli*, although it has been shown to be autorepressive in *Staphylococcus aureus*^39^. Using the titratable TF library strain for ZntR along with the *P*_*zntA*_ reporter plasmid from the Zaslaver library^40^, we are able to independently control both the co-factor and the TF copy number while quantitatively measuring *P*_*zntA*_ regulation.

**Figure 4.**
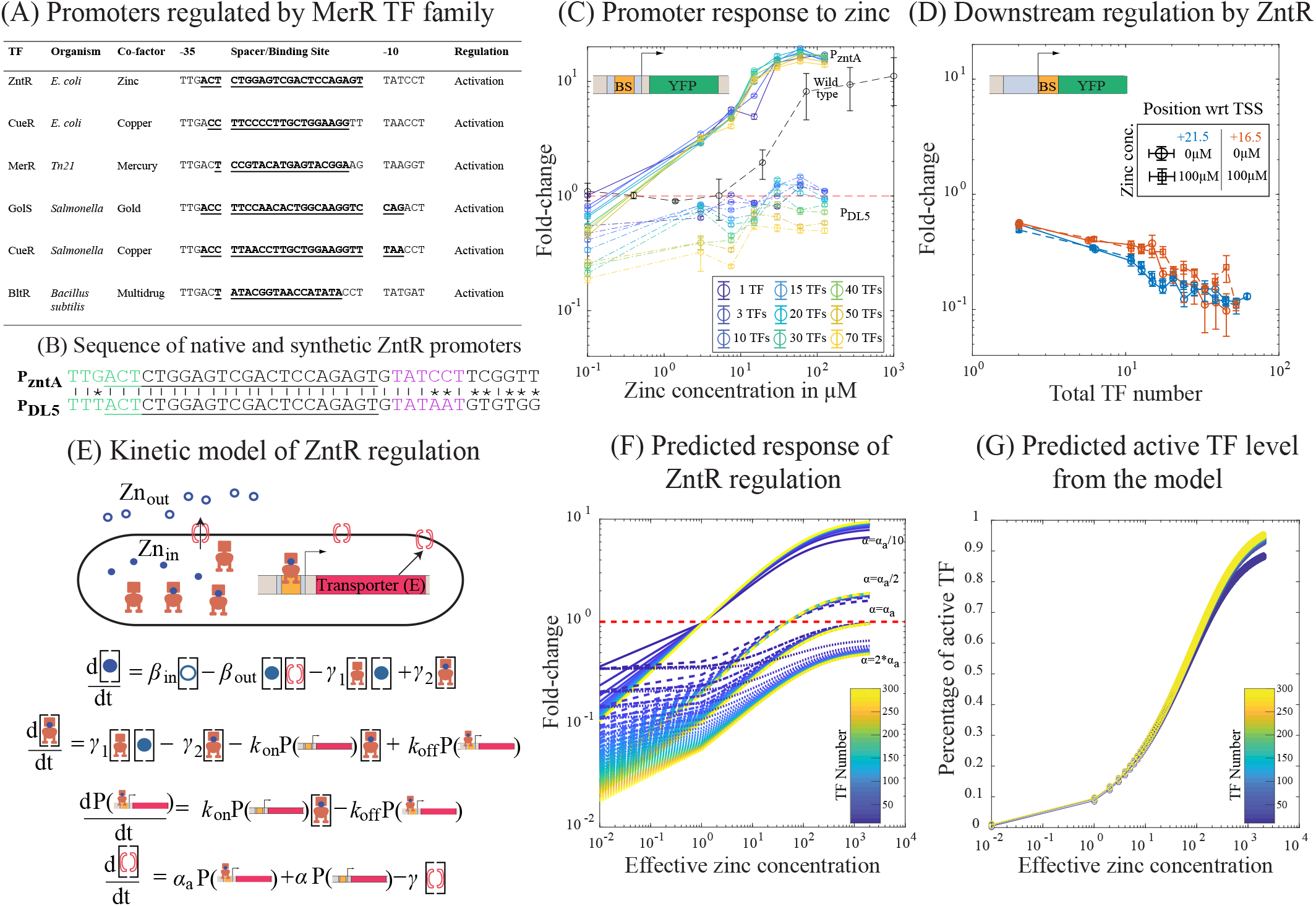
Regulation mediated by ZntR and Zinc: **(A)** Table showing the signature regulatory motif of the common MerR TF families in different bacteria. TFs of MerR family have their binding site acting as a spacer between the −35 and −10 boxes for RNAP binding. **(B)** Shows the sequence of *P*_zntA_ and *P*_DL5_ with ZntR binding site as the spacer. Shown in green is the − 35 element and in magenta is the − 10 element. The sequence of ZntR binding site in underlined. **(C)** Response curve for the *P*_zntA_ promoter in wild type (black dashed line) and in ZntR library strains (with fixed TF concentration (Solid line)) for different concentration of zinc. The dotted lines are the response of modified *P*_DL5_ promoter with ZntR binding site as the spacer. **(D)** Regulatory curves when the binding site for ZntR is downstream of the promoter (instead of being between the − 35 and − 10 boxes). The ZntR binding site acts as pure repressor independent of the zinc concentration when present outside the core promoter element **(E)** Simple kinetic model describing the regulatory features of ZntR-Zinc mediated regulation shown in Fig 4C. The reporter here is the transporter gene (E). *α* is the basal expression from the promoter for E and *α*_*a*_ is the acceleration brought about when the promoter for E is bound by an active TF (TF^*^ or Zinc-TF complex). **(F)** The output of the kinetic model when *α*_*a*_ is kept constant and the basal expression, *α* is altered. When *α* is less than *α*_*a*_ there is repression at lowest zinc concentrations that is dependent on TF copy number and activation at higher concentrations similar to the response form *P*_zntA_ promoter (solid lines in Fig 4C). When *α* is equal (dot-dash line) to or greater than (dotted line) *α*_*a*_ there is strong repression and weak activation and this regulatory feature is similar to that observed for modified *P*_DL5_ promoter with ZntR binding site (dotted line in Fig 4C). **(G)** Solving for active TF concentration indicates that the fraction of active TF (active TF/total TF) at a given zinc concentration is constant.

As described above, the *P*_*zntA*_ promoter contains the binding site for ZntR within the spacer sequence between the − 35 and − 10 box for RNAP binding (Fig 4B). ZntR uses zinc as co-factor although it also can weakly recognize other heavy metals like cadmium^41^. To understand the natural response function of ZntR, we measured the response of promoter *P*_*zntA*_ to different zinc concentrations in a standard laboratory strain (MG1655), as seen from the black dashed curve in Fig 4C. It is important to note that we only control zinc concentration in the media rather than the intracellular concentration, in practice the levels of internal zinc are complex, as the primary zinc exporters are changing with zinc and zinc is being bound by several metallo-regulatory proteins in the cell including RNAP^42^. However, the ZntA promoter clearly responds to the titration of external zinc and shows activation of transcription with a maximum fold-change of 10 at the highest zinc concentration when compared to the expression from a *zntR* knockout strain at the same zinc concentration. We next tested the effect of zinc titration on the same promoter construct in our library strain for several fixed TF copy numbers (solid color curves in Fig 4C). We see that ZntR has a zinc-dependent regulatory function that is not simply “inactive” to “active” but instead it changes from a repressor to an activator with zinc. Repression of the *P*_*zntA*_ promoter by ZntR can be small (roughly 10% or so) for low concentrations of TF and up to roughly 2 fold in the presence of hundreds of TFs. This repressive behavior was likely hidden in wildtype MG1655 by an innate feedback in ZntR regulation which would repress ZntR expression in the absence of zinc and thus prevent it from strongly repressing ZntA at low zinc. Our library strains are able to reveal this behavior due to the decoupling of natural regulation from the expression of the TF. Perhaps even more surprising is that activation of the promoter by the TF is largely independent of the total number of TFs; the color curves collapse for concentrations above roughly 1*μ*M of zinc. This means that the overall response of this system and the level of activation of exporter protein is robust to the expression level of the TF when zinc is present.

We wanted to further examine ZntR regulatory function by measuring regulation at different promoters and different locations. In Fig 4B, we show how we integrated the ZntR binding site as a spacer sequence into a common synthetic promoter derived from the *lac* operon, *DL5*^43^. Importantly, these promoters are very similar to the *P*_*zntA*_ promoter with only a single nucleotide change in the − 35 box and two changes in the − 10 box (see “asterisk” signs between the sequences in Fig 4B). The data is shown as dotted lines in Fig 4C for regulation of *DL5* by ZntR; we see slightly stronger repression for a given number of TFs which is alleviated by zinc. Importantly, we still see this dual function property, however in this case the activation is weak and does not outweigh the repressive function of the TF at any measured zinc and TF combination. As such, we find that the strength of both modes of regulation are sensitive to promoter sequence; in this case the repression was stronger and the activation was weaker than for ZntR acting on *P*_*zntA*_.

Next we measured ZntR regulatory function, both in the absence and presence of zinc, when binding at other locations on the promoter. We measured fold-change of the promoter with the ZntR binding site at two locations immediately downstream of the promoter (centered at +16.5 and +21.5). From previous work on a handful of TFs, we anticipate that TFs regulate at these locations through pure steric hindrance where the fold-change will be repressive and a reflection only of a single parameter, the occupancy of the TF^7,35,44–46^. In Fig 4D, we show the fold-change for these two binding locations as a function of TF copy number both without added zinc (solid line) and with 100 *μ*M zinc (dashed line). Significantly, the regulation both with and without zinc is the same which implies that activation by zinc does not alter the affinity of the TF for the binding site; this also suggests that the regulatory shift of ZntR with zinc at the native binding location emanates from a change in the regulatory function of the bound TF rather than a change in its occupancy. Fitting a simple steric hindrance model of regulation to this data enables measurement of the TF binding affinity^12^, which is the sole free parameter of the model. We find a binding affinity of − 13.8*k*_B_*T*, roughly equivalent in strength to LacI binding to the LacO2 operator^7^. Clearly the regulatory function of ZntR, even in this small number of examples, is incredibly flexible; naturally it is capable of providing zinc-dependent regulation that switches function from repression to activation depending on zinc but it also can serve as a TF insensitive to zinc availability.

To explore which features of the regulatory system enabled key features such as the robust TF number independent activation seen in Fig 4C, we built a simple kinetic model. The basic features of the model are highlighted in Fig 4E; expression of the target gene occurs at rate *α* in the absence of the TF. When the TF-zinc complex is bound, it acts as an activator increasing expression to a rate *α*_*a*_ *> α*. However, when zinc free TF is bound, it acts as a repressor capable of shutting off expression entirely. Zinc import is assumed to be proportional to the external concentration where as export is proportional to the concentration of ZntA in the cell. The results of Fig 4D allow us to estimate the on and off rates of TF binding of ZntR which we set equal regardless of TF-state (bound or unbound by zinc). We solve this model for steady-state and calculate the fold-change as a function of zinc concentration. We find (Fig 4F) that this model produces curves with a similar phenomena to the data in Fig 4C; at low zinc we see a copy number dependent repression, while with added zinc we see a collapse to copy number independent activation. The model enables us to probe the conformational state of the TFs in the cell to see how the collapse is achieved. As shown in Fig 4G, we find an important feature that the percent of TFs that are active is kept relatively constant independent of TF number in the cell.

### Case study 2: Regulatory role of paralogous TFs

The phenomenon of paralogous TFs, TFs within an organism that arise from gene duplication and divergence, is common throughout living systems^47,48^. Bacteria are particularly susceptible to this thanks for their propensity for lateral and horizontal gene transfer. Isorepressors are groups of TFs that share higher homology in the DNA-binding domain and form overlapping regulons such that they regulate identical (or similar) sequence motifs. Some examples from *E. coli* include pairs of TFs such as GadW and GadX as well as GalR and GalS but examples of isorepressor trios, also exist such as MarA, SoxS and Rob. Although these TF sets recognize similar consensus sequence motifs, their actual regulatory functions may differ at both quantitative and qualitative level.

Since our library strains can be extended to allow multiple single gene deletions, we aimed to measure the differences in regulation between isorepressors, GalR and GalS. To do this, we knocked out the paralogous TF in each of the GalR and GalS library strain (*i*.*e*. we knocked out GalR in the GalS library strain and vice versa). We then measured five native GalR-GalS responsive promoters in these strains at a range of induction levels. The native promoters have between 1 and 6 binding sites for GalR/GalS. The response of each of these promoters to titration of GalR or GalS is shown in Fig 5A. We find that four of these promoters have qualitatively similar responses to GalR and GalS, although the difference in magnitude of each response varies slightly with some promoters. On the other hand, the promoter for GalS (green curves, Fig 5A), is activated with GalR and repressed with GalS. Interestingly despite both TFs being categorized as repressors, we see significant levels of activation in response to GalR and GalS for some of these promoters especially, *P*_galP_ (yellow curves, Fig 5A). It is perhaps unclear from the data in Fig 5A if the difference in regulation arises from distinct TF function of each Gal paralog or if it is simply the result of differential affinity for the promoters.

**Figure 5.**
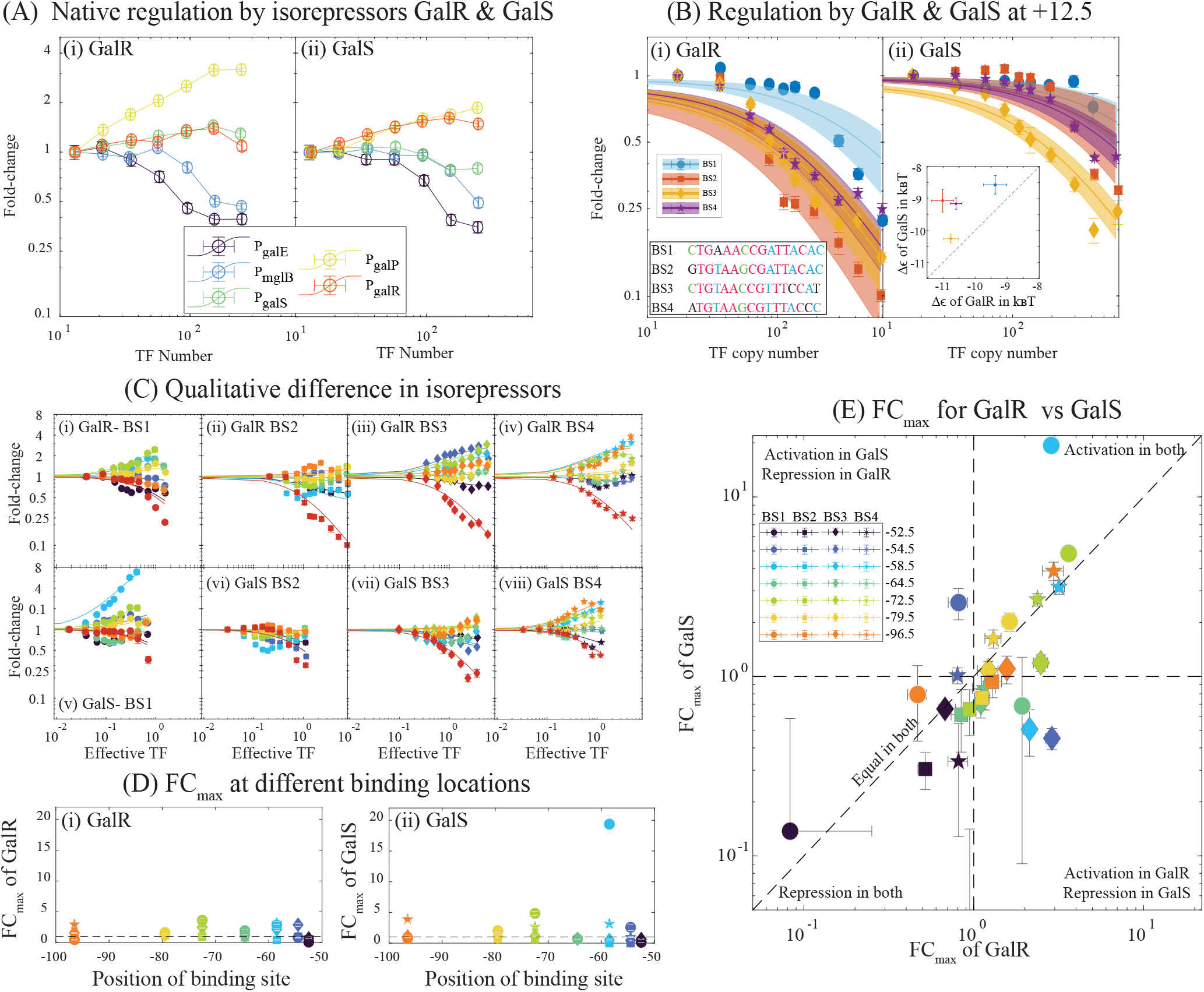
Quantitative and qualitative differences between the iso-repressors GalR and GalS: **(A)** Plot showing the response of GalR (i) and GalS (ii) to native promoters regulated by the isorepressors. **(B)** Regulatory curves for binding sites (BS1, BS2, BS3 and BS4) in GalR (i) and GalS (ii) at position +12.5 relative to the TSS. The insert in (i) shows the binding sequence of the 4 motifs analyzed in this study. Nucleotide common to all 4 binding sites are in red. Nucleotide found in at least 3 sequences are in blue and in at least 2 sequences are in green. Unique nucleotide are in black. The insert in (ii) shows the inferred binding affinity for different binding sites in GalR and GalS. **(C)** Analysis of the regulatory curves for BS1, BS2, BS3 and BS4 at different locations (− 52.5, − 54.5, − 58.5, − 64.5, − 72.5, − 79.5, − 96.5, *and* + 12.5) relative to the TSS indicate different qualitative features of the TFs at different binding location. **(D)** FCmax values at different binding locations for GalR (i) and GalS(ii). **(E)** shows a comparison of FCmax values of GalR and GalS at a given binding location. Data in each quadrant represents the unique features of GalR and GalS for the 4 binding sites.

We next tested four distinct binding site sequences; two sites that natively regulate *P*_galP_ (BS1 and BS2), one site from *P*_galS_ (BS3) and the last from *P*_galR_ (BS4) for regulatory differences in response to GalR or GalS. The natural location of these sites are at − 243.5 for BS1, − 61.5 for BS2 and BS3 and +8.5 for BS4. We measured fold-change for each individual GalR/GalS binding site placed at different locations along a synthetic promoter. In this case, the binding site we introduce represents the only known TF binding site on the promoter. Similar to our approach with ZntR, we first introduce each binding site directly downstream of the promoter. This data is shown in Fig 5B. The data for both TFs at each of the four binding sites fits well to the pure steric hindrance model of regulation allowing us to infer the binding energy of each sequence for either TF. The inset to Fig 5B compares these affinities and demonstrates that for each site GalR binds stronger than GalS. Interestingly, though, the rank order of sites is different for the two TFs; for both TFs BS1 is the weakest followed by BS4, however BS2 is the highest affinity site for GalR while BS3 is the highest affinity site for GalS. BS3 site has roughly equal affinity for GalR and GalS. These results indicate that the binding consensus for GalR and GalS are not the same.

Next, we infer the regulatory effect of GalR/GalS at other binding locations on the promoter. For this we use a general model of gene regulation that quantifies the regulatory role of the TF^35^,

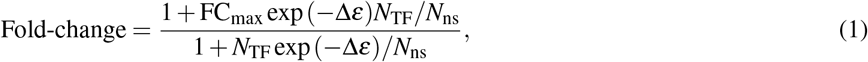

where FC_max_ represents the maximum fold-change at saturating TF concentration and exp(−Δ*ε*) is the affinity measured in the previous plots. Importantly, the steric model fit to that data is recovered by setting FC_max_ = 0. Fig 5C shows the fold-change as a function of TF copy number for both GalR and GalS binding to each of the four binding sequences at binding locations upstream ranging from − 52.5 to − 89.5. Each of these data sets fits well to the model with FC_max_, the regulatory function of the TF, as the sole fit parameter. Fig 5D(i,ii) show the regulatory role of each binding site for GalR (i) and GalS (ii) as a function of binding location. Surprisingly, we see several cases where one binding site at a position causes activation whereas a different sequence causes repression (*i*.*e* there are points both above and below the black dashed line at 1 in Fig 5D(i,ii)). Fig 5E compares the regulatory effect for GalR and GalS. Data point fall in one of the four quadrants of this plot, the upper right and bottom left quadrants are for binding sequences that have the same qualitative role for GalR and GalS regulation at that location. On the other hand, points in the top left and bottom right quadrants have different qualitative regulatory function for GalR and GalS; we see that the activation by GalR and repression by GalS is the most common form of differential regulation between the two TFs. Finally, data points that fall along the one-to-one dashed line have identical quantitative regulatory function with either GalR or GalS. Clearly despite having similar binding recognition, GalR and GalS can function as qualitatively different proteins beyond merely a preference for a given binding sequence and this difference depends on both binding sequence and binding location.

## Discussion

Genetic libraries have become an essential experimental resource in functional genomics. The primary goal of such genome-wide mutant collections is the unbiased study of all genes to reveal how each is involved in the dynamics and robustness of a given cellular process. In particular, there is a rich assortment of such libraries available in *E. coli* including a single-gene deletion library (the keio collection^20,21^), a transcriptional reporter library (the Zaslaver library^40^), and the ASKA open reading frame clones (the ASKA library^49^), among others. These mutant libraries have served as a stand-alone tool to address fascinating biological problems. For example, the keio deletion library has helped identify essential and non-essential genes across different growth conditions and facilitated the reconstruction of metabolic networks with higher precision^50,51^ and the transcriptional reporter library has been instrumental in obtaining the precision and dynamics in promoter activity under different extracellular perturbations^34^. Similar libraries have been constructed in other organisms, for instance single-gene deletion libraries exist for many bacteria such as *Bacillus subtilis*^52^, *Pseudomonas aeruginos*^53^, and *Acinetobacter baylyi*^54^ and also eukaryotic organisms such as *Saccharomyces cerevisiae*^55–57^. Importantly, these libraries have also played crucial roles in unexpected ways, for instance the keio library has been an essential tool in the study of modulating host-microbe interactions in dietary and drug responses of *C. elegans*^58–60^.

The titratable TF library introduced here is designed as a readily available genome-wide tool that enables quantitative control of individual TFs in *E. coli* (Fig 1A). Classic techniques for studying TF function rely on completely knocking out or over-expressing a given TF in order to infer the regulatory or physiology role. The resource introduced here provides the ability to control a TF in order to observe its role as a function of its concentration, which is measurable thanks to a fluorescent fusion to the TF. Circumventing natural TF copy number control is particularly important since TF genes are typically auto-regulating which makes TF control difficult; here the TF is expressed entirely from a synthetic promoter that can be induced with the small molecule aTC.

Many established tools, such as ChIP-seq and SELEX, have been developed for the purpose of determining where TFs bind and to which sequences they prefer to bind. Our library is designed to be a complimentary tool aimed at aiding studies seeking to determine a TFs regulatory function once bound (Fig 1A). The titratable TF library is versatile and can be combined with other single gene mutant library collections such as the transcriptional reporter library^40^, the keio single gene deletion library^20^ or the isolated regulatory position sweep library developed in our lab previously^35^ in order to make controlled measurements of TF regulation. The ability to isolate and control expression of individual TFs allows for characterization of the role of individual TFs without entanglement from their physiological function or role in higher order network effects such as autoregulation^8,61^. Similar strategies have previously enabled thorough characterization of individual TFs^7,12^, our library enables these approaches on a TF-wide level (Fig 1A).

We have demonstrated this use with two case studies examining regulation by the zinc-responsive TF, ZntR, and the paralogous TFs, GalR and GalS. In both cases we were able to isolate and quantify TF function through the use of the appropriate TF titration library strains. Another crucial tool in both of these studies is the presence of a quantitative model to interpret the data; the TF titration library enables characterization of gene regulation in a simplified system where TF number is under control which, in turn, enables simpler models with fewer parameters that are not intrinsic to the TF itself. For instance, in both examples we measured regulation of the studied TFs immediately downstream of the promoter as a way to infer the affinity of our TFs to a binding site, in one case this demonstrated that binding affinity of ZntR is independent of zinc concentration. Crucially for this case, knowing the affinity is not enough to predict regulation, you must also know the regulatory effect of the TF once bound. This is where our methodology of TF characterization shines.

In the other example presented here, we measure regulation for TF binding downstream of the promoter which demonstrated the differences in binding affinity of GalR and GalS to specific binding sequences. This is a crucial step towards disentangling affinity from function. As a result we could quantify how the TF function of the two paralogs differed in terms of not only their affinity for specific sequences but also in their function when bound to those sequences.

Immense progress has been made in genetic circuit design using TFs from a single family of TFs^9,62^. Overall, the goal of improving the repository of well-characterized TFs to include a greater diversity and more orthogonality has the potential to enable genetic circuit design of more complex responses with overall simpler architectures with fewer parts. Ideally, this library can be used to not only gain an understanding of natural regulation but to also improve our understanding of the general input-output function of TF regulation and expand the toolkit of synthetic biology (Fig 1A).

## Acknowledgments

We wish to thank Amir Mitchell, Job Dekker, Marian Walhout, and Michael Lee for helpful discussions. We declare no conflict of interest. Funding: Research reported in this publication was supported by NIGMS of the National Institutes of Health under award R35GM128797.

## Materials and Methods

### Construction of TF library

*E. coli* MG1655 is the parent strain used in our library construction. Single gene deletions of TF genes in the Keio library (with BW25113 as parent strain) is moved by P1 transduction into *E. coli* MG1655 expressing constitutive TetR at the *gspI* locus, and the kanamycin cassette associated with the keio knockout is flipped using the *frt* flippase expressed from the pCP20 plasmid. These strains serve as no fluorescence (and no TF) control strains for the corresponding titratable-TF strain in our library. Unless otherwise stated, all steps are performed in 96 well plates. Primers to amplify TFs are designed using customized matlab codes. Individual TF genes are cloned by Gibson assembly into pSC101 plasmid between the *P*_tet_ and aek-linker-mCherry sequence. Gibson clones were confirmed by sequencing and used as a template to amplify the *P*_tet_ -tf-aek-mCherry fusion gene for integration at the *ybcN* locus. Chromosomal integration is assisted by the lambda red recombinase proteins (*exo, beta*, and *gamma*) expressed from pKM208^63^ in wildtype MG1655. Successful integrants were confirmed by sequencing and moved by P1 transduction into the corresponding TF control strains.

### Growth Characteristics

For each experiment, 10 different TFs from the library (no TF control and the corresponding titratable TF strains) and wild type MG1655 are grown in 96-well plates to characterize the growth and mCherry fluorescence in M9-minimal media supplemented with glucose. The strains are grown overnight in LB and diluted 10^4^-fold into M9-minimal media with different aTC concentrations (0,1,3,5,7,9 and 15 ng/mL) and grown in 2mL volume 96-well plates at 37° C and 250rpm until it reaches OD600 of 0.4. 10 *μ*L of these cells are then diluted into 190 *μ*L of the same media in 300 *μ*L volume 96-well plates and transferred to plate reader with automated setting (TECAN MPro-200). OD600 values and mCherry fluorescence are measured every 30 minutes for up to 20 hours. Growth rate and steady-state mCherry fluorescence values are then calculated in the exponential phase of growth. Background subtracted OD600 values are log transformed and a polynomial fit is performed over a sliding window. A plot of the fit values across the sliding window gives characteristic regimes: a noisy regime for the lag phase, distinct peak in the growth phase and a plateau for the stationary phase. The maximum of the peak value corresponds to the growth rate of the particular strain. The growth rates of the test strains are normalized by the growth rate of MG1655 (grown in the same aTC concentration as the test strains) measured on the same day. Normalized growth rates are used in hierarchical clustering to determine different characteristic features of TF titration on growth rates. Complete linkage was used for the calculation of distance matrix. Pair wise correlation between TF number and growth rates are estimated using the inbuilt function (corrcoef) in MATLAB.

### Estimation of calibration factor

The calibration factor, *ν*, for the conversion of mCherry fluorescence to TF copy number is quantified as described in^43^, by measuring the stochastic fluctuations in fluorescence partitioning during cell division. Briefly, cells expressing the TF-mcherry fusion protein are grown as described in the section for Microscopy, and just before imaging 100 *μ*L of cells from different aTC concentrations are pooled together and washed twice with M9-glucose minimal media containing no aTC to stop any further production of mCherry. Cells are then spotted on 2 % low melting agarose pad made with M9-glucose minimal media. Phase images are captured for roughly 100 fields and their positions are saved for later. This phase images (named as Lineage tracker) will serve as a source file for lineage tracking of the daughter cell pairs (*I*_1_ and *I*_2_). After one doubling time (roughly 1 hour or depending on the doubling time for different TFs), the microscope stage was returned to the same field of view using the saved position matrix and are imaged again (and named as daughter finder) now using both phase and mCherry channels. For partition statistics to estimate the calibration factor cells saved as daughter finder are segmented using a modified version of Schnitzcells code^64^. Daughter pairs (*I*_1_ and *I*_2_) are picked manually by matching the segmented daughter finder with the phase image in lineage tracker and picking cells that made exactly one division. The mean pixel intensity and area of the daughter pairs are obtained using region props. The background fluorescence is estimated as described in^61^ using the inverse mask of individual frames. The sum and squared difference in fluorescence are estimated from the total fluorescence of the daughter cells after division. The resulting single cell measurements are binned for summed fluorescence values and fitted with a binomial distribution function to obtain the calibration factor *ν* for 10 different TFs used in this study. Only TFs that do not follow a volumetric partitioning during division can be counted by this technique and to do so the TFs are assumed to be dimerized and distributed randomly on the chromosome. Clearly as shown in Fig 1E, there are TFs (such as DhaR and NrdR) that are localized in the cell and might not follow random partitioning during division. Such TFs cannot be counted by this technique. 9 out of the 10 TFs measured had similar calibration factor and one TF, ArgR had higher value for *ν*. ArgR is hexameric protein and the DNA-binding is stabilized by the co-factor Arginine. Such factors might affect the estimation of protein number by stochastic fluctuation in fluorescence partitioning and hence, every TF should be carefully evaluated for the estimates of *ν*.

### Estimation of absolute TF numbers for the entire library

Since the mCherry levels at different aTC concentration for the library strain is measured in the plate reader, we needed to convert the measurements of TECAN to its microscopic equivalent before estimating for the absolute TF number. We measured 3 different TFs simultaneously in plate reader and microscope. We fit an exponential to the log-transformed values of the measurements from plate reader and the microscope in order to obtain the slope. We used this slope to convert all the data from the plate reader to its microscopic equivalent. We then used the mean of the calibration factor for 10 measured TFs to obtain the estimate for protein number at a given aTC concentration. The data from^23^ is used to compare the TF copies to the physiological concentration.

### Microscopy and Data Analysis

ZntR knockout strain (no-TF control strain) is used as an auto fluorescent strain for microscopic measurement and the same strain transformed with the corresponding reporter plasmid serves as the constitutive strain. The ZntR-TF-titration strain from the library is directly transformed with different ZntR reporter plasmids. For GalR-GalS reporter assays, the auto-fluorescence control strain is a double knockout of *galR* and *galS* and the constitutive strain is the double knockout strain transformed with the reporter plasmid. The titratable-TF-mCherry fusion construct for GalR and GalS from the library strain collection is P1-transduced into the double knockout control strain to make the corresponding titratable library strains. These strains are then transformed with the corresponding reporter plasmids for binding assay.

For microscopic analysis, control strains and their respective constitutive and titratable strains are grown overnight in LB media and diluted 10^5^-fold into fresh M9-minimal media supplemented with glucose and different concentration of aTC. These strains are grown for 16 − 18 hours until it reaches a OD600 of 0.2 − 0.3. For experiments with zinc as a co-factor, 0.1M stock of fresh zinc sulfate is added to the media and serially diluted to the desired zinc concentrations. As zinc sulfate is volatile the stocks are stored at − 20° C and thawed right before use. Unless otherwise specified, the aTC concentration used for all microscopic experiments are 0, 0.25, 0.5, 0.75, 1, 1.25, 1.5, 1.75, 2, 2.5, 3 ng/mL and 300*μ*L cells are grown in 2mL-deep well 96-well plate. Once in steady state, cells from different aTC concentration are pooled and washed twice with 1X PBS and spotted on 2 % low melting agarose pad made with 1X PBS. The constitutive strains and auto fluorescent strains are processed the same way as the pooled library strain and all samples are imaged on an automated fluorescent microscope (Nikon TI-E) with a heating chamber set at 37° C. For constitutive and auto-fluorescence sample 15 unique field of view are imaged resulting in roughly 100 − 300 cells per sample. For pooled titration strains, 40 unique fields are imaged per sample resulting in 1000 − 2000 cells per sample. ^64^

Segmentation of individual cells is performed using a modified version of the matlab code, Schnitzcells. We use this code to segment the phase images of each sample to identify single cells. Mean pixel intensities of YFP and mCherry signals are extracted from the segmented phase mask for each cell using region props, an inbuilt function in matlab. The auto fluorescence is calculated by averaging the mean intensity of the auto fluorescence strain in both mCherry and YFP channel and is subtracted from each measured YFP or mCherry values. Total fluorescence for each channel is obtained by multiplying the mean pixel-intensity with the area of the cell. Fold-change in expression for a given binding site is calculated by the ratio of total fluorescence of strains expressing the TF to the strains with no TF. Fold-change is also calculated for the TF strains to the knockout expressing target YFP constitutively, without any regulation by the TF. The final fold-change is the ratio of fold-change with regulation and without the regulation. mCherry values are converted to TF number using the measured calibration factor for each individual strain. The values are binned for TF number.

### Kinetic model for ZntR regulation

We built a simple kinetic model to explore the regulatory features of ZntR (TF) mediated regulation of ZntA transporter (E). Based on Fig 4D, we assume that the binding affinity (*k*_on_, and *k*_off_) is the same for both TF and TF^*^ (zinc bound TF). For simplicity the transporter (E) itself is the reporter gene here. The basal expression from *P*_off_ is dictated by rate *α* and *α*_*a*_ is the rate when the promoter is co-driven by polymerase and TF^*^.

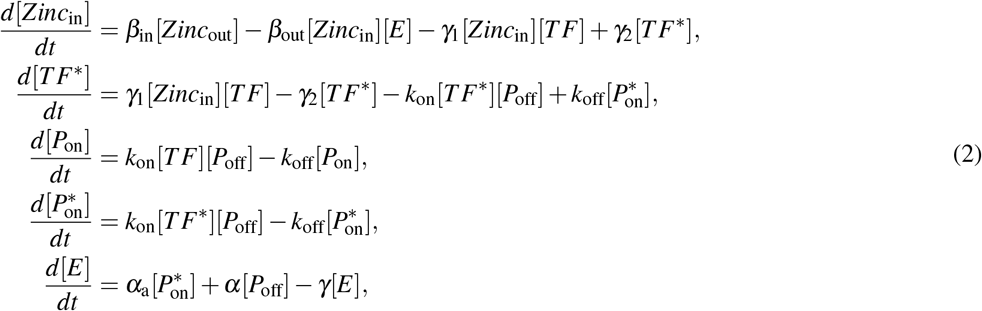

Here, *Zinc*_in_ and *Zinc*_out_ are the extra and intracellular concentrations of Zinc and *β*_in_ and *β*_out_ are their corresponding rates. *TF* and *TF*^*^ are the concentrations of the free TF and zinc-bound TF, respectively. Both TF and TF^*^ binds DNA with a rate *k*_on_ and unbinds with a rate *k*_off_. *γ*_1_ and *γ*_2_ are the rates at which the TF binds and unbinds intracellular zinc. *P*_on_ and 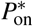 are the concentrations of promoters bound by TF and TF^*^. Only TF^*^ bound promoters 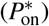 will initiate transcription with a rate of *α*_*a*_. *γ* is the degradation rate of the transporter, E. Constraints on the model are 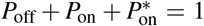 and 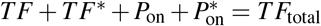. The right hand side of the above equations is set to zero in order to obtain the steady state values of each component.

### Supplementary Figures

**Figure S1.**
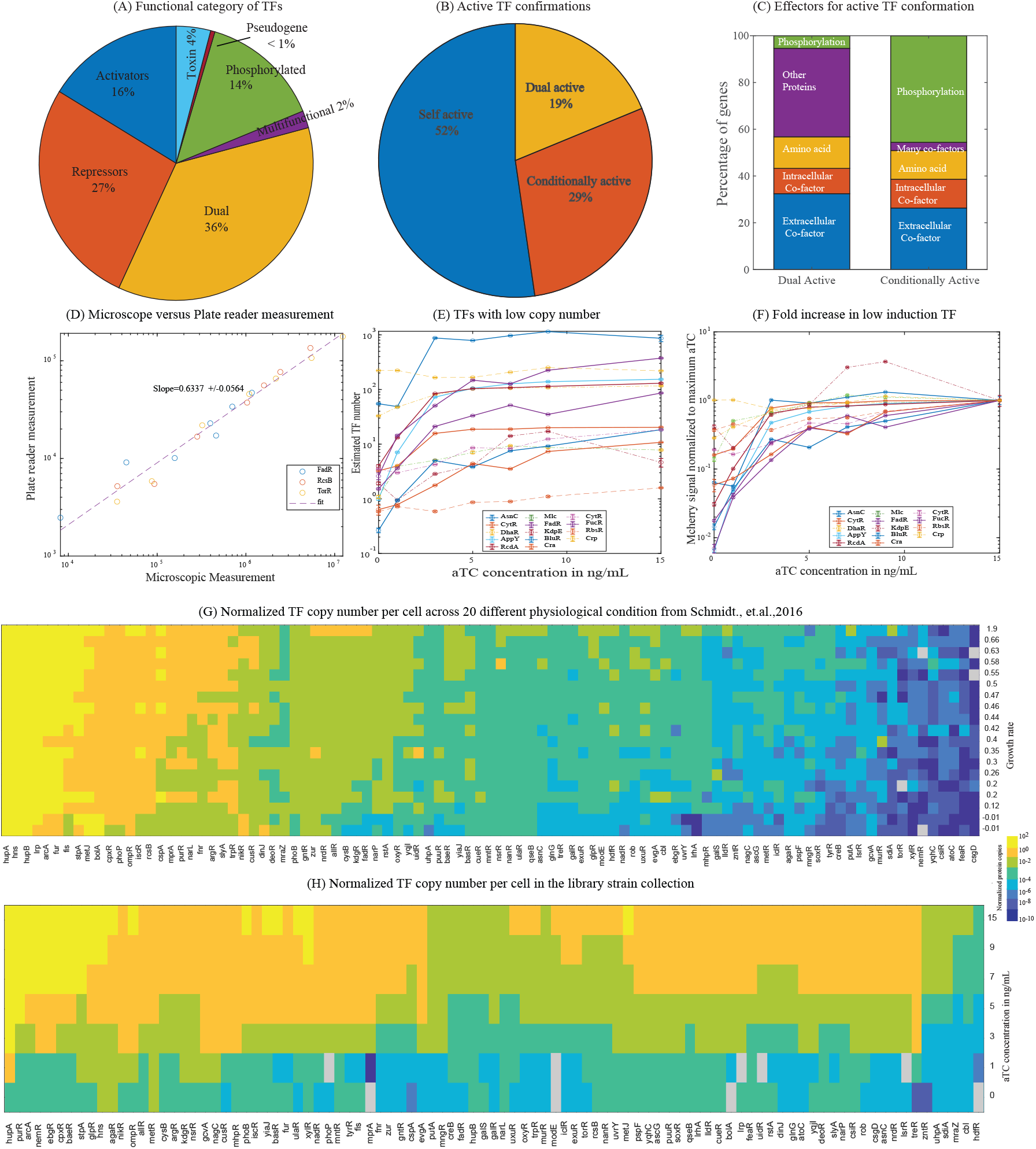
Transcription factor library: **(A)** Based on the mode of action, TFs can be classified into seven functional categories: Activators, repressors, dual regulators, toxin-antitoxins, phosphorylated TFs, pseudogenes and multi-functional TFs. **(B)** TFs can be classified into 3 categories based on the conformation with which they bind the DNA *i*.*e*. active TF conformation. **(C)** The effector molecules used by TFs to facilitate DNA binding or active TF conformation. **(D)** Conversion of measurements from the plate reader to the measurements from the microscope. **(E)** Measurement of mCherry fluorescence of selected strains (that showed no-titration in TECAN measurements) in microscope. **(F)** Data in Fig S1E represented as normalized to maximum aTC concentration. TFs shown in dashed lines shows no titration. **(G)** Physiological concentration of different TFs (normalized to the average TF obtained in the library strain at maximum induction) in wild type strains as measured by Schmidt *et. al*.^23^. As shown in this figure, altering the growth rate is not a good proxy to achieve different levels of TFs due to the inherent regulations of the native TF. **(H)** Every TF in our library strain is titratable to 100-1000s of TF copy number under identical induction condition (normalized to the average TF obtained in the library strain at maximum induction).

**Table S1.** Kinetic rates used in the simulations for Fig 4F

| Rates | Symbols | Value |
| --- | --- | --- |
| Degradation rate | $\gamma$ | $0.1 \text{ s}^{-1}$ |
| Binding energy<br>(Experimentally verified) | $k_{\text{on}}/k_{\text{off}} = \exp(-\Delta\epsilon)/N_{\text{ns}}$ | 0.2 |
| Rate of<br>constitutive expression | $\alpha$ | $5\text{s}^{-1}$<br>$10\text{s}^{-1}$<br>$1\text{s}^{-1}$<br>$20\text{s}^{-1}$ |
| Rate of activation | $\alpha_{\text{a}}$ | $10 \text{ s}^{-1}$ |
| Effective zinc concentration | $Zinc_{\text{out}}(\gamma_1/\gamma_2)(\beta_{\text{in}}/\beta_{\text{out}})$ | $0.01 - 2000\mu M$ |

**Figure S2.**
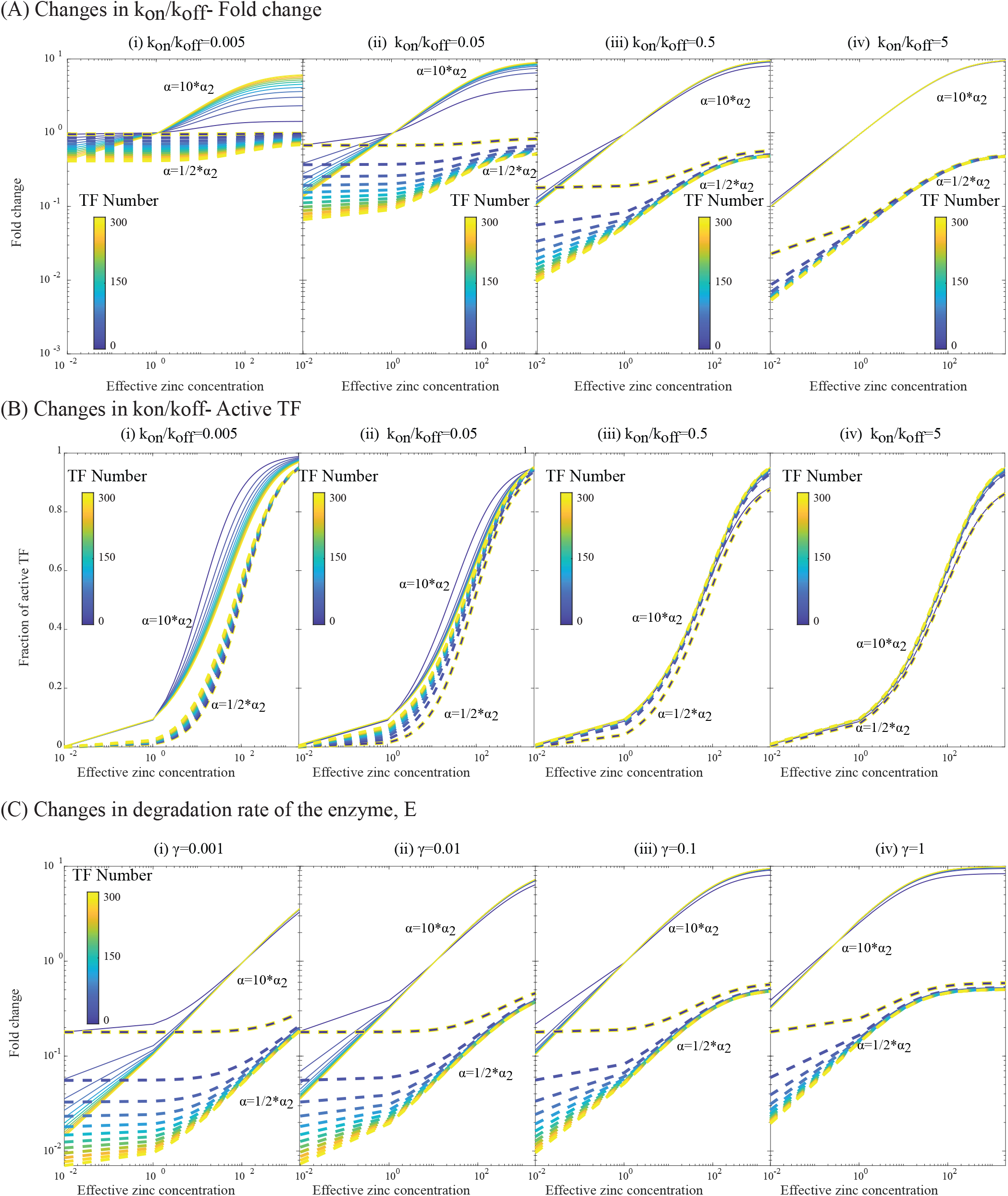
Model parameters: **(A)** *k*_on_ is the binding rate per free TF per unit time, *k*_off_ is the unbinding rate per unit time. The ratio of *k*_on_*/k*_off_ is the kinetic equivalent of binding energy of the TF. A small *k*_on_*/k*_off_ ratio indicates weaker binding (and vice versa).

**Figure S3.**
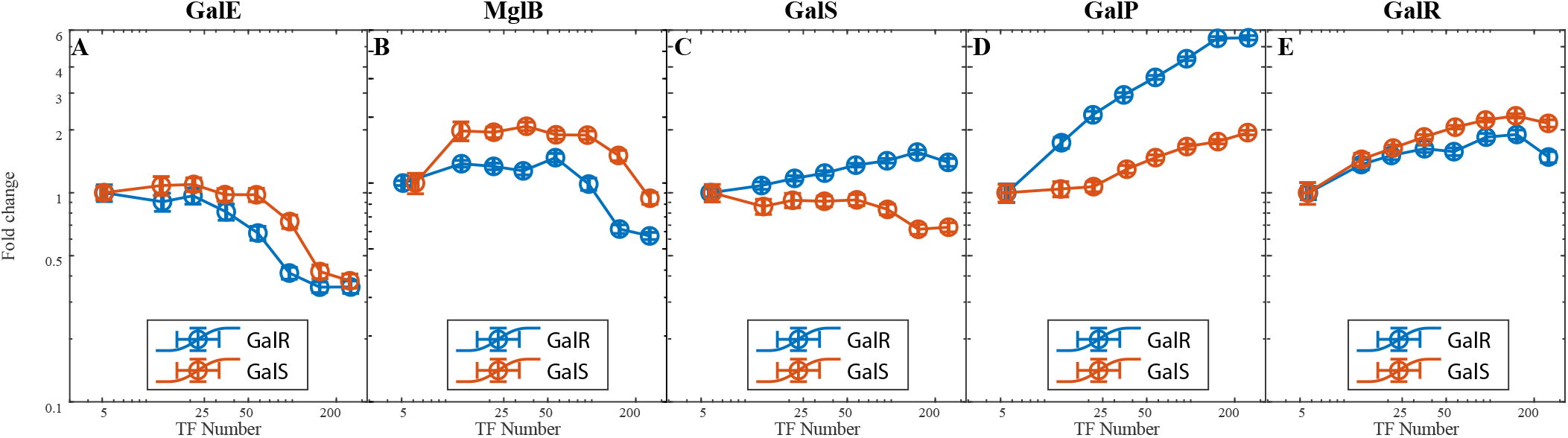
Isorepressors on their native promoters: **(A-E)** The regulatory curves for GalR and GalS on their native promoters from the Zaslaver transcriptional reporter library.

### Measurements of calibration factor

Calibration factor is a measure of stochastic event and care should be taken to avoid or minimize contributions from extrinsic factors during experimental set up and data processing. Major factors affecting the experimental set up are (1) efficient washing of cells in order to remove any traces of aTC and shut-down any further expression of mCherry, (2) incubator associated with the microscope should be equilibrated to 37° C several hours before the experiment, (3) the precision in the time to capture one division (images taken too early might result in far less cells and images taken several minutes after the first division might have volumetric difference), and (4) finally the lower signals (major contributors of the y-intercept or the calibration factor) is largely dependent on efficiency with which background signals are calculated. As we have shown previously^61^, background fluctuations are critical especially for lower signals that are just couple of counts above the backgorund. Hence, we use error-propagation to account for any errors that could arise from changes in background counts in estimating the sum of the fluorescence or in squared difference in fluorescence.

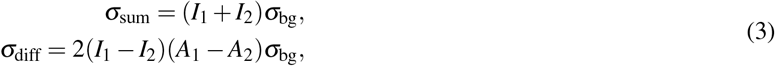

Here, *σ*_sum_, *andσ*_diff_*arethestandarddeviationinthemeasureo f sumanddi f f erencein f luorescencebetweenthedaughtercellsandσ*_bg_ is the standard deviation of the auto-fluorescent strain. *σ*_bg_ is measured experimentally. *σ*_sum_, *andσ*_diff_ are inferred from the above relationship. A *σ*_bg_ value of upto 5 is tolerable in the measurements of calibration factor (Fig S4C). The day-to-day variability in the measurements of calibration factor for one of the library strains is given in (Fig S4A). Roughly two fold difference is observed between different days for the measurement of *ν*.

**Figure S4.**
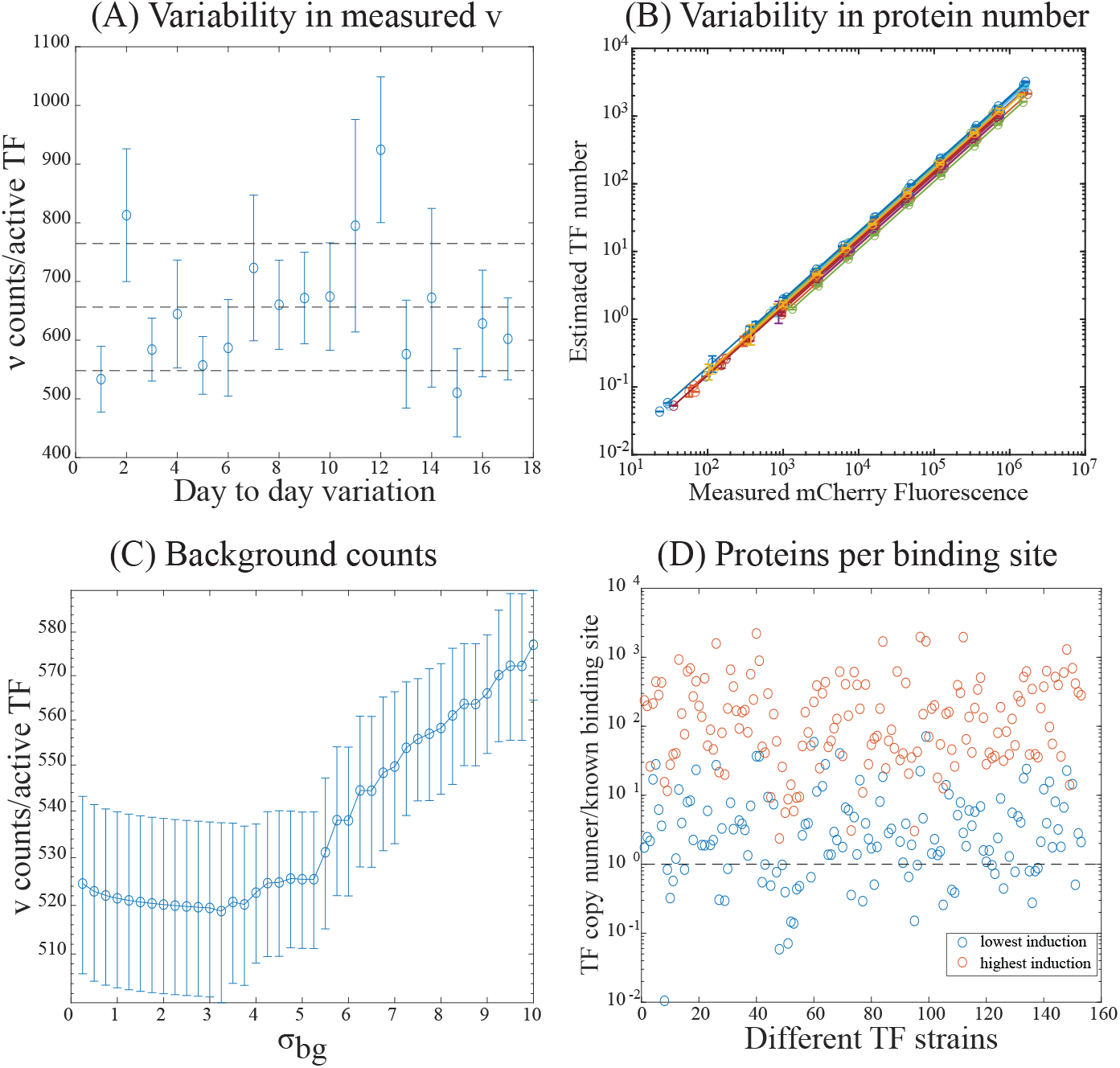
Measurement of protein copy number: **(A)**Plot showing the measured value of *ν* for a single TF over multiple days. **(B)** Even when the measured *ν* values are varied over multiple days as shown in (A), the TF copy number across different days stays fairly constant.**(C)** Plot showing the impact of fluctuations in the background values in estimating *ν*. **(D)** Plot showing the predicted TF copy number at lowest and highest induction normalized to the corresponding TF’s total known binding site reported in regulonDB. For most TF’s even at the lowest induction condition there is at least 1 TF per known binding site

### Phosphorylated Transcription Factors

The response regulators of the two component signal transduction system of *E. coli* are the TFs that require phosphorylation and there are a total of 26 of them in our library strain collection. These TFs present additional challenges in quantitative studies as these proteins take up distinct regulatory roles proportional to its phosphorylation status inside the cell. Discrete protein kinase, usually encoded along side the associated response regulator (or the TF) is essential for phosphorylation of the corresponding TF. However, in our library strain, only the response regulators’ concentration is controlled and not the concentration of kinase. In addition, the kinase is usually encoded within the same operon expressing the response regulator and in our library strains the response regulators are deleted in-frame from its native locus. Such deletions could easily alter the expression of the kinase associated in the same operon. Hence, phosphorylated TFs might need additional genetic modifications such as deletion or regulated expression of the regulated kinase or replacing the critical amino acids of response regulator with phospho-mimetic amino acids^65^ or choice of growth characteristics that would naturally induce phosphorylation. Although we have succeeded characterizing the phosphorylated TF, CpxR^35^ in the past, for most other TFs phosphorylation remains challenging as in S5. CpxR might be special case as it activates the expression of its own kinase but may not be generalized for other kinase and need careful characterization.

**Figure S5.**
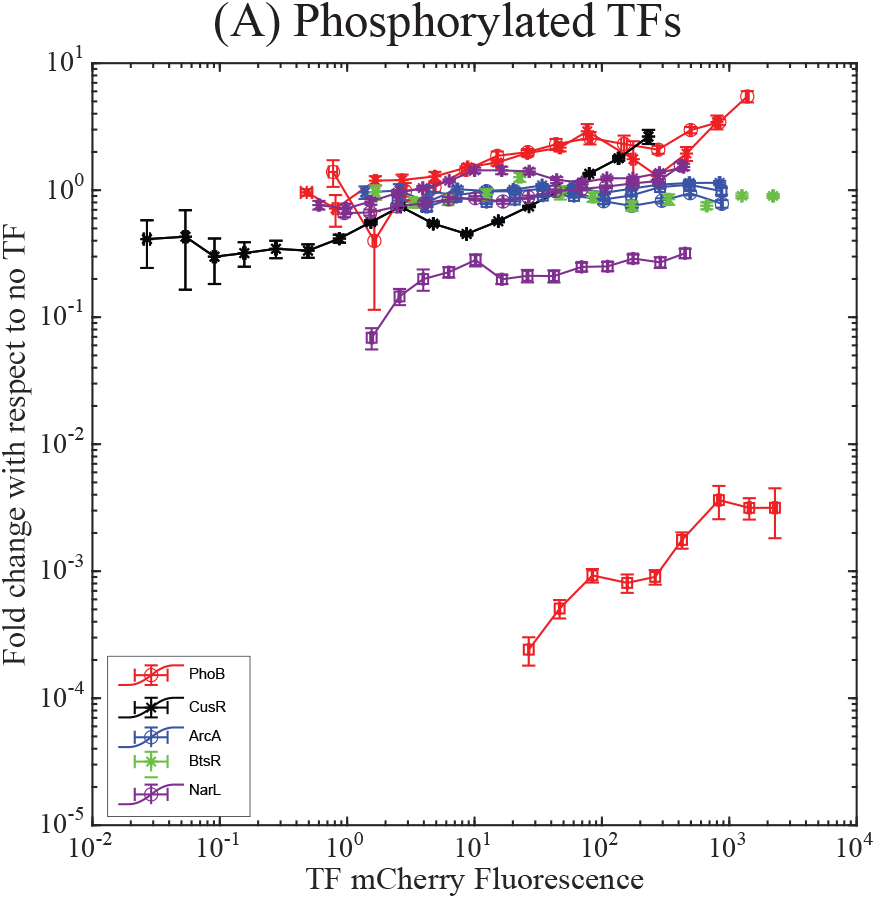
Input-output relationship for selected phosphorylated TFs. **(A)** Input-output relationship is measured for TFs: PhoB, CusR, ArcA, BtsR and NarL with selected plasmids from Zaslaver’s transcriptional reporters. These group of proteins require phosphorylation for its response regulation.

### Global regulator H-NS

Global regulator, H-NS is one of those few TFs whose predicted TF concentration is lower than the reported physiological concentration (Fig 2E). About 5% of genes in *E. coli* is regulated by H-NS^66^. Hence, we tested the physiological effect of titration of H-NS in several minimal media (Fig S6A-B). As shown in panel B. the mCherry levels are similar in different media (M9-glucose, M9-arabinose and M9-glycerol media) however, the amount of aTC required to rescue growth equivalent to the wild-type is different under the 3 tested condition with growth in glycerol requiring more H-NS proteins to rescue wild-type like growth than glucose or arabinose minimal media. This also demonstrates yet another functionality of our library strains .i.e. in studying the limitations of TF titration under different nutrient conditions.

**Figure S6.**
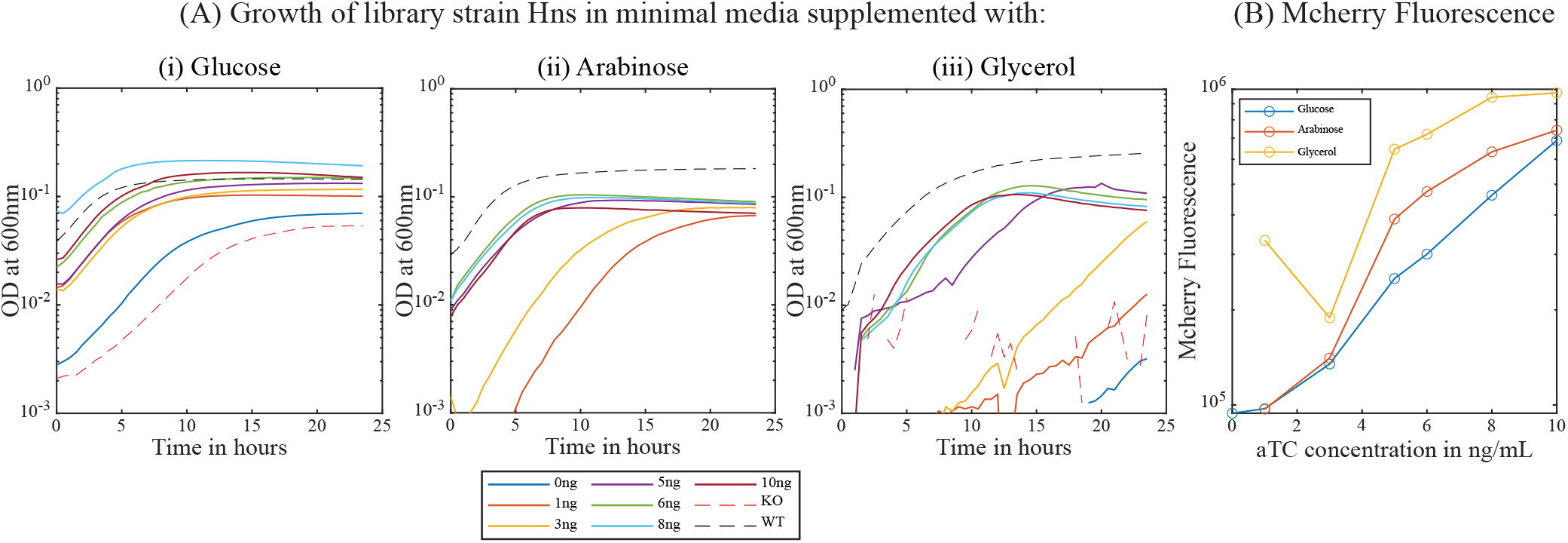
Global regulator H-NS. **(A)** Plot showing the growth of H-NS knockout and titratable library strains in minimal media supplemented with (i) Glucose (ii) Arabinose or (iii) Glycerol as a sole carbon source. H-NS is a global regulator and one of those library strain’s with the TF numbers lower than the physiological concentration. **(B)** Plot of the mCherry levels as a function of aTC concentration in different growth medias for the library strain, H-NS.

### Physiology of TF titration

**Figure S7.**
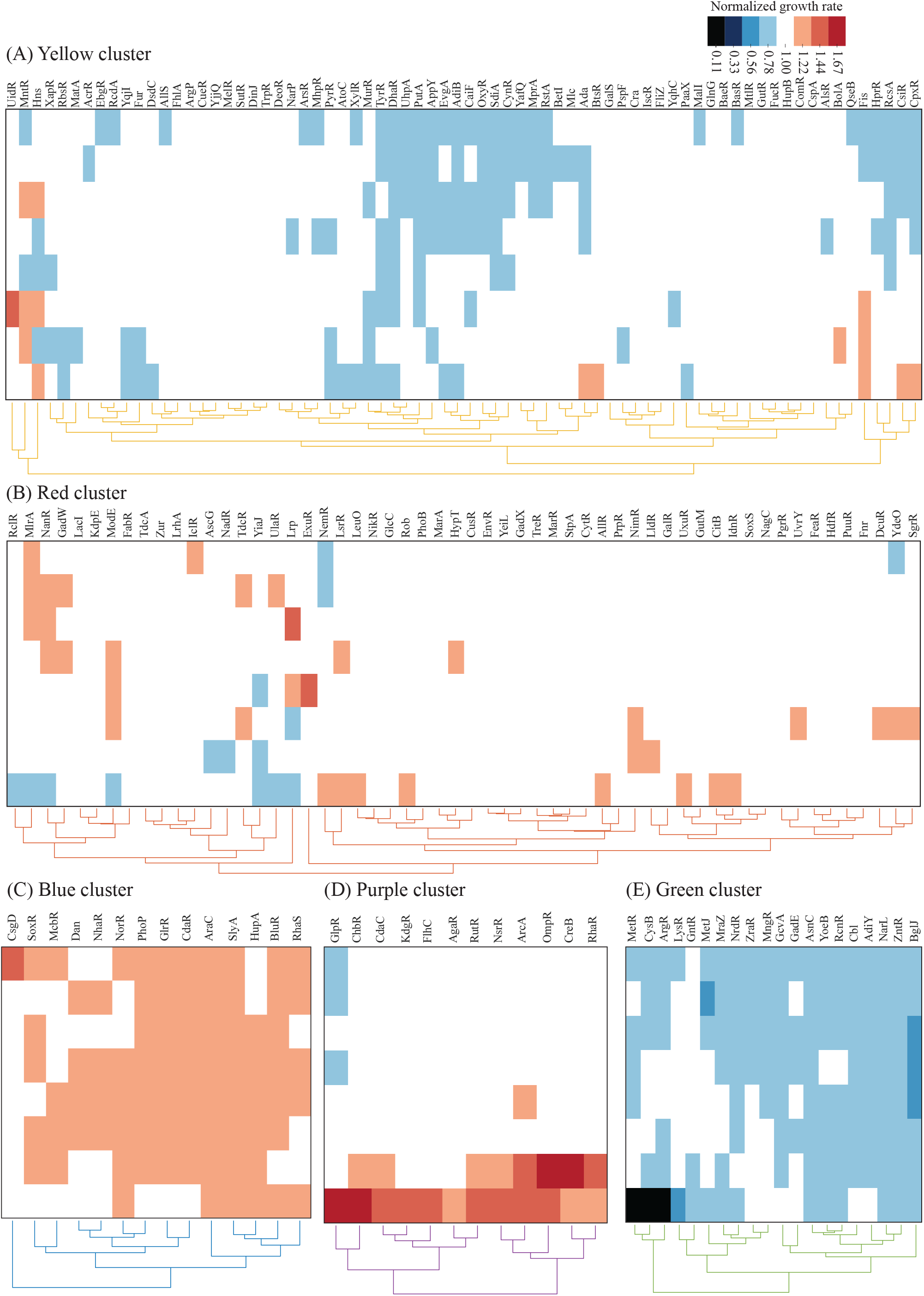
Individual clusters: Data in Fig 3A with 5 individual clusters presented separately and gene names displayed. The last cluster in Fig 3A includes only 2 TFs (Nac and PdhR) and is not shown here.

